# Improvements to the rice genome annotation through large-scale analysis of RNA-Seq and proteomics datasets

**DOI:** 10.1101/300426

**Authors:** Zhe Ren, Da Qi, Nina Pugh, Kai Li, Bo Wen, Ruo Zhou, Shaohang Xu, Siqi Liu, Andrew R Jones

## Abstract

Rice (*Oryza sativa*) is one of the most important worldwide crops. The genome has been available for over 10 years and has undergone several rounds of annotation. We created a comprehensive database of transcripts from 29 public RNA sequencing datasets, officially predicted genes from Ensembl plants, and common contaminants in which to search for protein-level evidence. We re-analysed nine publicly accessible rice proteomics datasets. In total, we identified 420K peptide spectrum matches from 47K peptides and 8,187 protein groups. 4168 peptides were initially classed as putative novel peptides (not matching official genes). Following a strict filtration scheme to rule out other possible explanations, we discovered 1,584 high confidence novel peptides. The novel peptides were clustered into 692 genomic loci where our results suggest annotation improvements. 80% of the novel peptides had an ortholog match in the curated protein sequence set from at least one other plant species. For the peptides clustering in intergenic regions (and thus potentially new genes), 101 loci were identified, for which 43 had a high-confidence hit for a protein domain. Our results can be displayed as tracks on the Ensembl genome or other browsers supporting Track Hubs, to support re-annotation of the rice genome.

## Introduction

The development of next-generation and third generation sequencing technologies mean that genome sequences are now being routinely generated for an ever expanding range of species, strains, breeds, and even individuals within populations. For the genome to be useful for fundamental and applied research requires high-quality annotation. Following genome assembly, annotation involves the discovery of the start codons of all genes, and their exon splicing patterns, which is a highly challenging task. Gene finding in most genome projects is performed via software that makes *ab initio* predictions of coding sequences, and where possible uses homology to other annotated genomes. Experimental data in the form of large scale RNA Sequencing (RNA-Seq) is also commonly used to find mRNAs and align reads that cross intron junctions to infer splicing. Undoubtedly the use of large scale RNA-Seq data vastly improves genome annotation but nevertheless, all genomes suffer from some proportion of mistaken annotation, such as incorrect translation initiation sites, incorrect splicing or pseudogenes called as protein-coding.

It is now becoming widely recognised that inference of the protein-coding elements of the genome can be greatly improved using large-scale mass spectrometry (MS) data on peptide sequences, in so-called *proteogenomics* approaches (1). In a typical proteogenomics approach, MS/MS spectra are searched against a customised protein sequence database, produced from curated gene predictions, as well as predicted possible sequences from *ab initio* gene finders and/or aligned RNA-Seq derived transcripts. Therefore, proteogenomics not only provides expression level evidence of protein-coding genes but also has the potential to improve the protein-coding gene sets by providing evidence that novel transcripts or alternative predictions for known genes have supporting evidence at the protein sequence level. There have been several proteogenomics studies on plants that have shown the ability to discover novel protein-coding genes and predict or improve splicing annotation. For instance, in 2009, Castellana *et al.* performed a proteogenomics analysis on *Arabidopsis* tissues (2). They successfully identified 778 novel genes and made 695 gene model refinements. Later in 2014, they developed an automatic method of proteogenomics and performed analysis on *Zea mays*, finding 165 novel protein-coding genes and proposing updated models for 741 additional genes (3).

Rice (*Oryza sativa*) is the staple food for half the world’s population. With the completion of the rice genome, it has become possible to elucidate the layers of information encoded by the sequence (4–6). Comprehensive genomic and transcriptomic studies of rice have been conducted worldwide, serving as a base for research aimed at matching the demand of increasing food supplies (7–12). Herein, we have performed a proteogenomics analysis on rice through collecting public genomics, transcriptomics and proteomics data, to discover novel protein-coding genes and new splice sites.

With the development of genomics, transcriptomics and proteomics techniques, the ability to detect ever higher proportions of the transcribed genes and evidence for translated proteins has become possible via proteogenomics. In addition, tools and strategies, such as customProDB and SpliceDB (13), have effectively improved the performance of proteogenomics by facilitating improved design of the search database. The construction of “novel event” candidates (i.e. new exons or splice junctions) is one of the most important steps in proteogenomics studies. Some studies aim to be comprehensive, such as using six frame translation of whole genome sequence (14,15), while these approaches are likely to contain exonic sequences for all possible genes, they suffer from a lack of statistical power (due to overall database size) and splicing information. An alternative is to use *ab initio* gene predictions from gene finding software (16). In this case, the constructed candidates will contain predicted splice event, but rely upon the accuracy of gene finding software which is not generally high, meaning that some possible splice sites will be missed. A third alternative is to create a search database from mapping RNA-Seq data onto the genome. The use of RNA-Seq results can balance these aspects, keeping a relative comprehensive search space but without the size expansion of six frame translations.

We designed our proteogenomics pipeline as follows. First, for database construction, we used transcriptomics data aligned onto the genome. For the translation step, we kept only the longest frame to control the overall database size. Second, for database searching we used multiple search engines via our previously published IPeak approach (17). IPeak combines the machine learning approach of Percolator and the FDRScore algorithm for search engine integration, which has been demonstrated to improve sensitivity over using a single search engine (18). IPeak is available as part of the mzid Library and ProteoAnnotator projects (19,20). Third, we performed extensive filtration to ensure that identified peptides not matching the official annotation (novel peptides) were high confidence and the corresponding spectra could not be explained by other causes. Fourth, to validate and annotate the resulting novel peptides and corresponding novel events, we aligned our novel peptides back onto the genome for visualisation against other tracks of evidence. Last, to standardise the presentation of results, we use standard formats from the Proteomics Standards Initiative (PSI) – mzIdentML (21) and proBed (22), which allow for rapid and automated visualisation of the results via public genome browsers. Through the use of 29 datasets of RNA-Seq data (789,141,453 reads), and 9 MS/MS datasets (2,051,418 spectra), this study represents one of the most comprehensive proteogenomics efforts undertaken on rice.

## EXPERIMENTAL PROCEDURES

An overview of the pipeline used for rice proteogenomics is summarised in Figure 1. The workflow is mainly divided to two parts for the processing of the RNA-Seq and MS/MS data, as follows.

**Figure 1.**
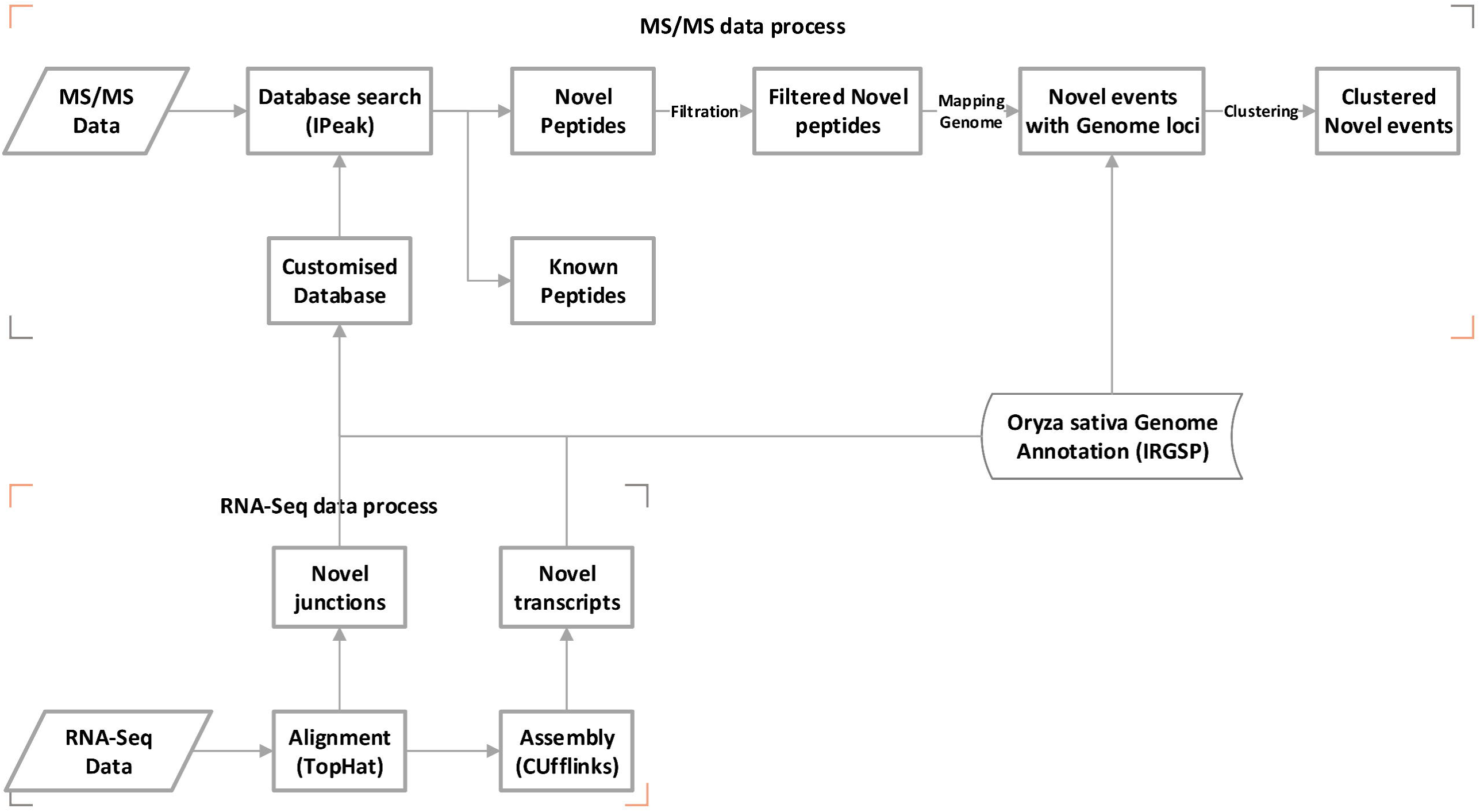
An overview of the analysis workflow for proteogenomics in this study.

### Data Collection

In this study, raw RNA-Seq data that was generated from the Illumina platform in paired end mode was collected from the European Nucleotide Archive (ENA, http://www.ebi.ac.uk/ena) database. A total of 29 runs (153,907,936,648 bases/789,141,453 reads) was contained in the datasets, and the full details are listed in Supplementary Table 1 (within SupplementaryResults.docx).

To search for peptide evidence, MS/MS data acquired from high-resolution mass spectrometers (LTQ Orbitrap XL, LTQ Orbitrap Velos, TripleTOF 5600 and Q Exactive) was used, regardless of whether the data was generated from profiling or enrichment studies. The raw MS/MS data for this study was collected from the ProteomeXchange (PX, http://www.proteomexchange.org/) database, including a total of nine datasets and 2,051,418 MS/MS spectra. Detailed information about datasets is listed in Table 1.

**Table 1.** The raw MS/MS data collected for this study from ProteomeXchange database.

| Dataset Identifier | Title | Instrument | Publication | Announce Date | Spectra count |
| --- | --- | --- | --- | --- | --- |
| PXD000265 | Oryza sativa egg, sperm, callus, pollen and seedling proteome | LTQ Orbitrap XL;<br>LTQ Orbitrap XL | Abiko *et al.* 2013 (30) | 2013/8/2 | 253743 |
| PXD000313 | Quantitative proteomics study on rice embryo during embryogenesis by using isobaric tags for relative and absolute quantification (iTRAQ) | Q Exactive | Zi *et al.* 2013 (31) | 2013/8/6 | 976822 |
| PXD000923 | Rice Pistil LC-MSMS | TripleTOF 5600 | Wang *et al.* 2014 (32) | 2014/7/31 | 87502 |
| PXD001030 | Proteomic analysis of proteins related to rice grain chalkiness using iTRAQ based on a notched-belly mutant with white-belly | TripleTOF 5600 | Lin *et al.* 2014 (33) | 2014/6/17 | 301027 |
| PXD001058 | Unravelling the proteomic profile of rice meiocytes during early meiosis | LTQ Orbitrap Velos | Collado-Romero, Alós & Prieto 2014 (34) | 2014/8/13 | 92162 |
| PXD002291 | Rice (Oryza sativa) lysine-acetylation LC-MS/MS | Q Exactive | Xiong *et al.* 2016 (35) | 2016/2/24 | 94400 |
| PXD002739 | Acetylome analyses in the germinating rice seed | Q Exactive | He *et al.* 2016 (36) | 2016/1/26 | 34344 |
| PXD002740 | Succinylome analyses in the germinating rice seed | Q Exactive | He *et al.* 2016 (36) | 2016/1/26 | 25862 |
| PXD003156 | Gel-free/label-free proteomic analysis of developing rice grains under heat stress | LTQ Orbitrap | Timabud *et al.* 2015 (37) | 2015/12/17 | 185556 |

### Construction of the customised database based on RNA-Seq data

The RNA-Seq reads from each run were individually aligned using TopHat (v2.0.12) against the *Oryza sativa* genome (IRGSP-1.0.30). The accepted matches in Bam format and the junctions in BED format were produced by TopHat. All the accepted reads from each run were individually sent for assembly into transcript sequences by Cufflinks (v2.2.1). Afterwards, Cuffmerge was employed to combine the transcripts from each run to form longer transcripts in GTF format. The longer transcripts marked with class code “=” are from the transcripts completely matched to the known exons, termed as *known transcripts*, while those with other class codes are from the transcripts partially or totally mismatched to the known exons (IRGSP-1.0.30), assigned as *novel transcripts (NTs).* All the *novel transcripts* in GTF format were taken for construction of the customised database. All the junctions were first de-duplicated and aligned against the “official junction sites” from the *Oryza sativa* annotation (IRGSP-1.0.30) to filter out the *known junctions* by custom scripts, and the remaining junctions were considered as *novel junctions (NJs)* for further construction of the customised database. All the novel junctions and novel transcripts are collectively called *novel events (NEs)*.

All the *NEs* were matched back to their corresponding genome loci. The matched genomic fragments were translated in six reading frames. An accepted novel translation product from six reading frame translation was judged by two criteria, more than 5 amino acids (15 nucleotides) at least, and only the longest product being taken for a transcript. All the accepted novel translation products were added in to the list of the *Oryza sativa* proteins annotated from IRGSP-1.0.30, as well as contaminants from cRAP (http://www.thegpm.org/crap/), to generate a new protein database for MS/MS searching.

### Peptide search based on MS/MS data (IPeak search)

IPeak, a Java-based open source software package, was employed for peptide search, which uses the Percolator to re-score peptide-spectrum matches (PSMs) from MS-GF+ (v9733), MyriMatch (v2.2.8634) and X! Tandem (v2009.10.01.1). IPeak incorporates the FDRScore algorithm to combine the results from different search engines. All the MS/MS data collected from nine datasets were converted into MGF format using ProteoWizard (v3.0.4238) (23), then were searched with IPeak against the customised database. The PSMs with FDRScores less 0.01, corresponding to q-value (global FDR) < 0.01, were initially used to create the list of peptides identified. Most search parameters in the original publications associated with each of the nine datasets were used in the IPeak search. All the identified peptides through IPeak derived from the known rice proteins were marked as *known peptides*, while that not from those proteins were denoted as *putative novel peptides (PNPs)*.

### Mapping *PNPs* to genomic positions

The *PNPs* were mapped back to the genome to locate their positions on the chromosome by custom scripts to generate the GTF files, which record the genomic positions of the corresponding *NEs* of *PNPs*. The positional information of *PNPs* and *NEs* were imported into the original mzIdentML file using the proteogenomics encoding described in the mzIdentML version 1.2 specifications (24).

### Filtration to remove the PNPs with potentially incorrect assignments or low confidence

The *PNPs* obtained from the IPeak search were further filtered to remove the potential for alternative explanations of the corresponding spectra being more likely. PEAKS PTM (25) is capable of identifying many types of post-translational modifications (PTMs) and chemical modifications to peptides. To reduce the possibility of misidentification of novel peptides due to modifications (e.g. where a novel peptide has the same mass as a known peptide with a common modification), the MS/MS spectra that were matched to the novel peptides were re-searched with PEAKS PTM against the official annotation. For any novel peptide whose spectrum was confidently identified by PEAKS PTM as the modified or alternative form of a known peptide (q-value < 0.01), was flagged for removal from the novel peptide set.

To further check that PNP identified sequences could not be explained by other types of biochemical events (missed cleavage, proteolysis or single amino acid substitutions) on known peptides, all the *PNPs* were aligned by BLASTp and custom scripts (using a regular expression) against the known protein sequences (IRGSP-1.0.30). The BLASTp search was conducted in two modes, default and short sequence optimised, with results filtered allowing for a maximum of one mismatch, zero gap, and exact sequence length between query and hit. Any *PNPs* with a confident match by BLASTp (either mode) were excluded from the result set. To further increase the confidence of *PNPs*, the peptides with only a single spectrum support were also removed. After all these filtration steps, the remained *PNPs* are called the *final novel peptides (FNPs)*.

### Clustering of FNPs and NEs on the genomic landscape

The *FNPs* and their related *NEs* were parsed from mzIdentML into GFF format by custom scripts, and were input to BEDTools (v2.25) to cluster the NEs onto the corresponding genomic loci. During clustering, 100bp was set as the maximum distance allowed between the *NEs*.

### Homology analysis of the FNPs with other plant proteins

To provide further evidence supporting the annotation of *novel peptides*, the sequences of FNPs were aligned using BLASTp (short sequence optimised mode to all the plant proteomes stored in Ensembl plants (release-38) with tolerance of only one residue mismatched or deleted in BLASTp.

All the transcripts corresponding to peptides located within intergenic region, and thus potentially new genes, were first translated to amino acid sequences, then the translation products were searched with InterProScan to identify potential protein domains, using filtration of e-value < 10^-5^.

### Visualisation of the NEs and FNPs

To visualize the *NEs* and *FNPs*, phpMs (26) was employed to transfer mzIdentML to proBed format, which is compatible with genome browsers supporting BED, such as IGV, UCSC, and Ensembl. To allow downstream comparisons, the two proBed files were combined from nine proteomics datasets: one contains all novel peptides identified and the other contains all peptides (non-novel) detected in the known *Oryza sativa* protein database. Track hubs are web-accessible directories of omics (genomic and proteomic in our project) data that can be viewed alongside official genome annotation through Genome Browser interfaces. Following the steps in https://genome.ucsc.edu/goldenpath/help/hgTrackHubHelp.html#Setup, one hub (containing two tracks) was generated, by converting the proBed files into BigBed format. The hub is hosted on our server and made publicly accessible via the track hub registry (http://trackhubregistry.org/ Search for “rice proteogenomic”). Each novel peptide and its cluster has its own link to Gramene/Ensembl browser which can be found in SupplementaryFile1.xlsx (tab “Final Novel Peptides”).

### Coding potential estimation

The coding potential (CP) of identified novel events clusters was estimated by CPAT (27). Rice known coding transcripts (42,132) and noncoding transcripts (total 53,250) from IRGSP-1.0.30 were used for training following the guide in (http://rna-cpat.sourceforge.net/). The three features, ORF size, Fickett score and Hexamer score, were taken to assess the training result. The training and thresholding for CPAT is exemplified in Supplementary Figures 4 and 5.

### ProteomeXchange submission

Raw files from all nine proteomic datasets and our result mzIdentML v1.2 files have been deposited to ProteomeXchange Consortium via the PRIDE (28) partner repository with the dataset identifier PXD008960.

## RESULTS

### Defining the novel events based on RNA-Seq data

In IRGSP-1.0.30, there are 91,080 genes with 97,751 annotated transcripts, including 35,679 protein coding genes, with 42,132 potentially protein coding transcripts. Within the annotation there are 150,594 annotated *official junction sites* of which over 98% of the sites (147,696) are contributed by the coding transcripts. The RNA-Seq data collected for this study from 29 runs exhibited an average mapping rate to the genome of 78.99%, with an average coverage to the annotated transcripts of 86.56% and to the (protein) coding transcripts of 97.48% (Supplementary Table 2 and Supplementary Figure 1). High coverage of the RNA-Seq data demonstrated the customise database was a relatively comprehensive collection of coding production of *Oryza sativa*.

The RNA-Seq data was aligned to the genome by TopHat. As a result, a total of 2,940,788 junctions (355,323 junction sites) were identified. By comparing with the *official junction sites*, 1,047,488 junctions (56,432 non-redundant junction sites) in the RNA-Seq data exactly matched to those annotated junctions, while the remaining 1,893,300 junctions (298,891 junction sites) were marked as *NJs*. Details of all the junctions and the corresponding sites are shown in Supplementary Table 3. The mapped reads were sent to assembly, leading to 201,360 transcripts constructed by Cuffmerge, in which 103,707 transcripts were exactly matched with annotated transcripts and the other 106,653 transcripts were regarded as *NTs.* Details of the assembled transcripts in each run are shown in Supplementary Figure 2. Thus, the *NE* set consists of 1,997,007 events.

A six frame translation of a genome sequence is a common method of looking for novel coding regions. However, a six frame translation of the IRGSP-1.0.30 genome generates 25,761,390 small ORF candidates using the length cut-off of 6 or more amino acids, which would be an excessively large search space, leading to reduced statistical power. On the bases of the customised database derived from RNA-Seq data described above, only about 1/26 small ORF candidates (970,204) were generated with the same length cut-off, 869,872 from *novel junctions* and 100,332 from *novel transcripts.* Further comparison of the theoretical digestion with trypsin (without missed cleavage), generates 31,088,508 peptides in a standard six frame translation database compared to 2,045,553 in the customised database, the search space for the latter being only 1/15 of the former. As such, we can conclude that the assembled customised database should allow for a much higher statistical power than a standard six frame translation.

### Identification of the novel peptides based on the collected MS/MS datasets

Through IPeak search at 1% PSM FDR against a total 9 ProteomeXchange datasets, 421,913 peptide spectra matches (PSMs) were found, corresponding to 47,663 peptides identified. The detailed statistics for the identified spectra and peptide are shown in Table 2. The PSM rates in these datasets were diverse, with generally higher rates in the datasets from enrichment studies, whereas lower rates in the datasets from profiling studies. Of the matched peptides, 43,495 peptides were derived from proteins in the IRGSP-1.0.30 annotation assigned as *known peptides*, corresponding to 8,187 protein groups, while the other 4,168 peptides were only mapped to the NEs, then termed as *putative novel peptides (PNPs)*. The details of PNPs are listed in Supplementary File 1 (tab “Putative Novel Peptides”).

**Table 2.** The overall counts of spectra, PSMs, percent of spectra identified, total identified peptides and total novel peptides per dataset.

| ProteomeXchange<br>ID | Spectra<br>Count | Total<br>Identified<br>Spectra<br>(FDR 1%) | Total Spectra<br>Identification<br>Rate<br>(FDR 1%) | Total<br>Identified<br>Peptides<br>(FDR 1%) | Putative<br>Novel<br>Peptides<br>(FDR 1%) |
| --- | --- | --- | --- | --- | --- |
| PXD000265 | 253743 | 83083 | 32.74% | 21563 | 1500 |
| PXD000313 | 976822 | 72516 | 7.42% | 17802 | 1182 |
| PXD000923 | 87502 | 16584 | 18.95% | 6344 | 705 |
| PXD001030 | 301027 | 29902 | 9.93% | 7678 | 466 |
| PXD001058 | 92162 | 24634 | 26.73% | 9458 | 605 |
| PXD002291 | 94400 | 16471 | 17.45% | 2654 | 172 |
| PXD002739 | 34344 | 9151 | 26.65% | 2913 | 173 |
| PXD002740 | 25862 | 10683 | 41.31% | 2934 | 196 |
| PXD003156 | 185556 | 158889 | 85.63% | 8213 | 314 |
| Total | 2,051,418 | 421913 | 20.57% | 47663 | 4168 |

The PSMs for *PNPs* and *known peptides* we refer to as nPSMs and kPSMs, respectively. Although we took the global FDR to control the false discovery rate of total identified PSMs, the nPSMs might suffer from a higher false positive rate than the global FDR estimated due to the method of database construction (29). To confirm the confidence of the nPSMs, we used three search engines, X!Tandem, MS-GF+ and MyriMatch, and attempted to address whether the distribution of Posterior Error Probabilities (PEP) for kPSMs and nPSMs was different from that generated from the decoy database. As shown in Figure 2A, the PEPs of kPSMs and nPSMs are evenly distributed along the three axes with the similar patterns and are scattered away from the origin, whereas the distribution of PEPs of decoy PSMs is completely different from that of kPSMs and nPSMs, and is narrowly located around the origin. Using the FDRScore to integrate search engine scores, Figure 2B reveals that the distribution of -log (FDRScore) of the nPSMs is comparable with that of kPSMs, but is significantly different from that of decoy PSMs. The matched ions, including b ions, y ions and b-y ion pairs, of nPSMs are compared with those from kPSMs as presented in Figure 2C. Although the right tail of the distribution of matched ions and ion pairs of kPSMs is somewhat higher than that of kPSMs, the distribution apex and shape are comparable. Overall, the evaluation illustrated in Figure 2 provides support that the MS/MS quality of the nPSMs are similar to kPSMs.

**Figure 2.**
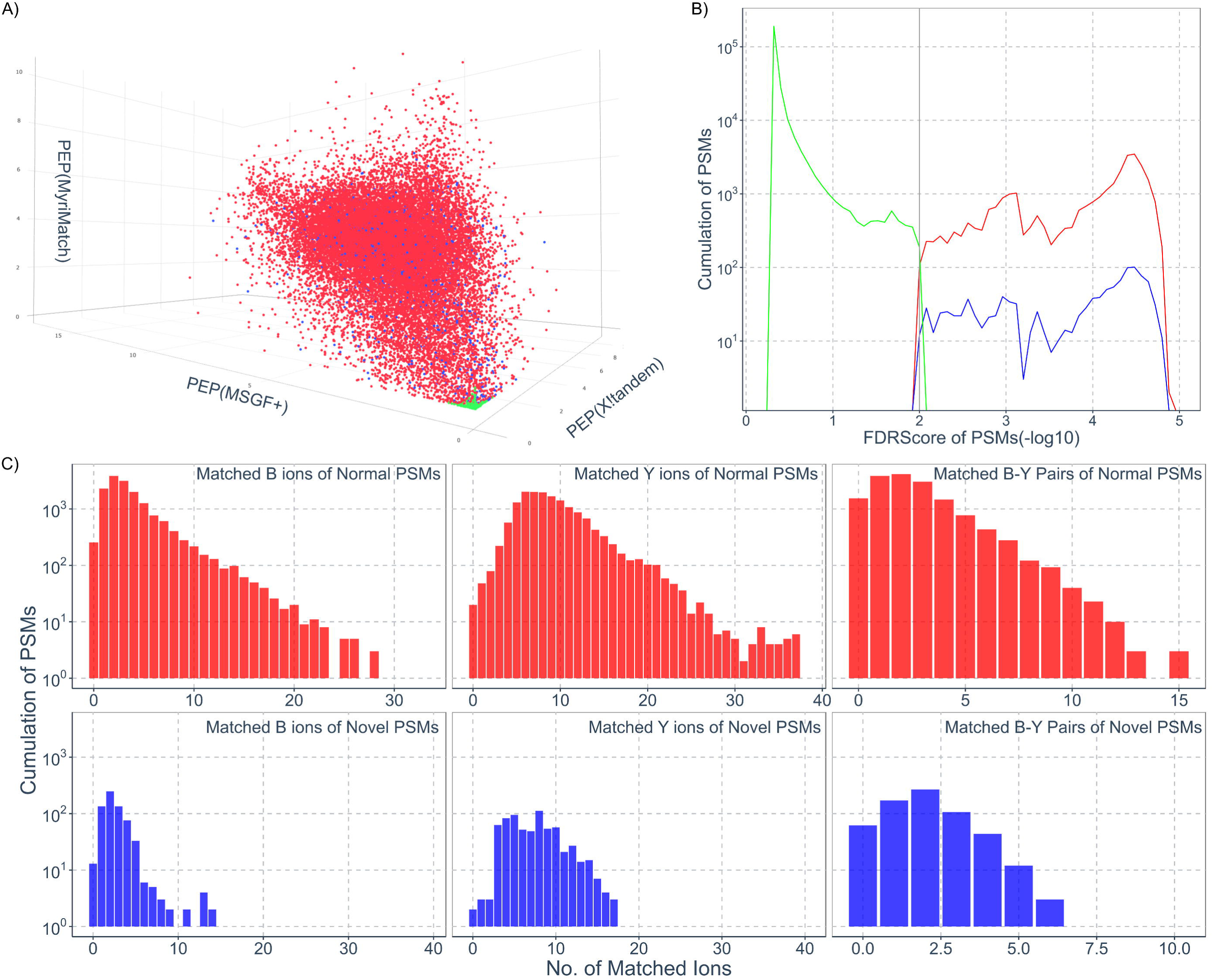
Evaluation of the novel peptides derived from MS/MS data. **A)** The overall scoring integration towards the novel peptides identified by three search engines. The axis in the three-dimensional scatter plot stand for –log 10 (Posterior Error Probability) values from XlTandem, MS-GF+ and MyriMatch, respectively. In the figure, the colour points represent the normal peptides (red), the novel peptides (blue) and the excluded peptides including decoy peptides and peptides with FDRScore over 1% (green), respectively. **B)** The distribution of FDRScores for the peptides with density plots. The colour curves have the same meaning as described in A). **C)** The distribution of matched ions for b, y and b-y pairs with bar plots. The colours represent the same meaning as described in A) (ions from excluded peptides are not shown).

### Filtration to remove the PNPs with potentially incorrect assignments or low confidence

As mentioned in Experimental Procedures, we filtered PNPs by searching their spectra with PEAKS PTM to search for evidence of known peptides with modifications or amino acid substitutions potentially explaining the same spectra. The 14,164 spectra corresponding to 4,168 PNPs were searched with PEAKS PTM, resulting in 740 spectra matching 169 peptides from canonical rice proteins with single amino acid mutation or common modifications at 1% FDR (Supplementary File 1, tab “PEAKS PTM Identifications”). To further ensure the PNP sequences could not have been derived from known proteins, we performed several filtration processes, regular expression matching, standard BLASTp and short sequence optimized BLASTp. First, a regular expression matching was conducted using custom scripts. A total of 403 peptides were matched by regular expression matching, including 397 peptides matched with a peptide in the official database generated without tryptic termini. Second, a BLASTp search with at most 1 mismatch was conducted to remove the peptides with potential single amino acid variations. Using standard BLASTp and short sequence optimized BLASTp, 542 and 674 peptides were matched. Combining all the filtration results, a total 904 distinct peptides were filtered out, which were matched by at least one of the filtration conditions. Figure 3A contains a Venn diagram displaying the detailed information regarding the filtration results. Figure 3B displays the peptides filtered in relation to the number of spectra supporting the identification, showing in most cases that there were few differences across different matched sets. One observation from Figure 3B is that the PEAKS PTM filter remove a relatively large proportion of peptides with >6 spectra support, likely indicating that these were otherwise confidently identified abundant peptides. For Figure 3A & B, we conclude that the different filtration mechanisms are complementary, since no single method was able to identify all possible causes of potential incorrect assignment.

**Figure 3.**
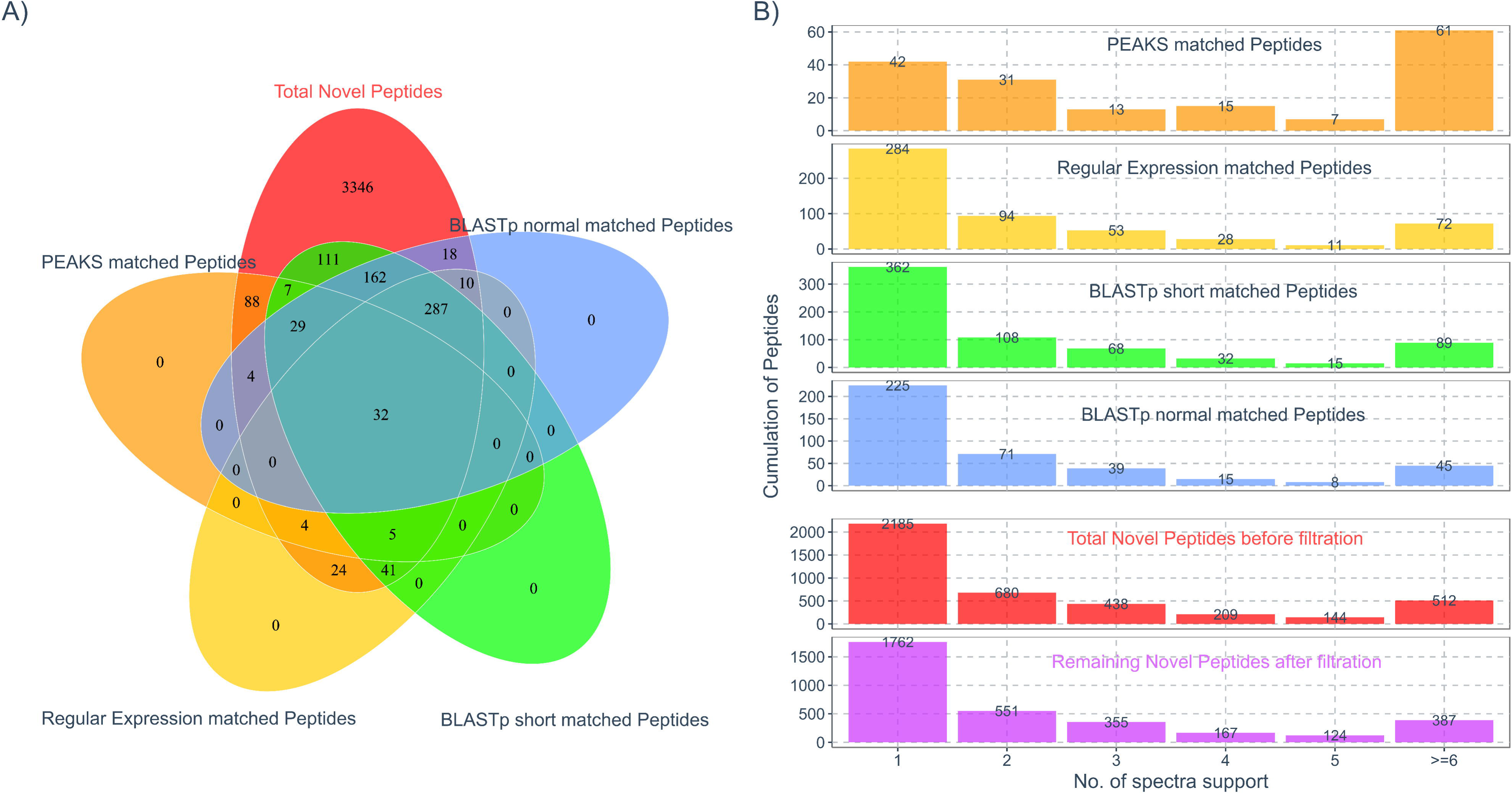
Multiple filtration to treat the putative novel peptides. **A)** Venn diagram indicating the overlap results of multiple filtration schemes for the putative novel peptides. The filtration includes PEAKS PTM (orange), BLASTp short search (yellow), BLASTp normal search (green) and regular expression match (blue). After the filtration, a total of 3,346 of the putative novel peptides (red) remain. **B)** The spectrum/peptide distribution passing and removed by filtration. The upper panel (four blocks): the colour bars indicate the distribution of spectra and peptides removed by the different filtration methods as in 3A. Lower panel (two blocks): comparison of the distribution of spectra and peptides for the total novel peptides with and without filtration.

After these filtrations, we found a large portion of the PNPs still supported by only a single spectrum (bottom panel of Figure 3B). In order to increase the confidence of PNPs, we adopted a more stringent criterion that an identified peptide must be supported by at least two MS/MS spectra. Thus, a total 1,762 peptide were further removed, leaving 1,584 peptides passing all filters, and thus marked Final Novel Peptides (*FNPs*, see Supplementary File 1, tab “Final Novel Peptides”).

### Clustering of FNPs and NEs on the genomic landscape

FNPs and their corresponding NEs were mapped back to genomic coordinates. Compared with IRGSP-1.0.30, 68 FNPs were mapped to multiple genome loci, while the other 1,514 were mapped to unique genome loci, called unique mapping FNPs. Among the 1,514 unique mapping FNPs, 944 novel peptides were located in intergenic regions, while the other 570 novel peptides are located in intragenic regions e.g. within introns, different frames or different splices of existing genes. However, the relatively large number of intergenic peptides does not necessarily indicate entirely new genes being discovered, since the matching transcripts might be located close to existing genes, and thus be additional or alternative exons. This point is further addressed by the clustering process described below. The details of the FNPs mapped to the rice genome are summarized in Table 3. In addition, 682 out of 1514 unique mapping FNPs span multiple exons potential junction sites. Importantly, 62 FNPs have also since been independently confirmed as protein-coding by the up-to-date annotation in IRGSP-1.0.38, implying that our analysis for discovery of novel peptides is valuable.

**Table 3.**
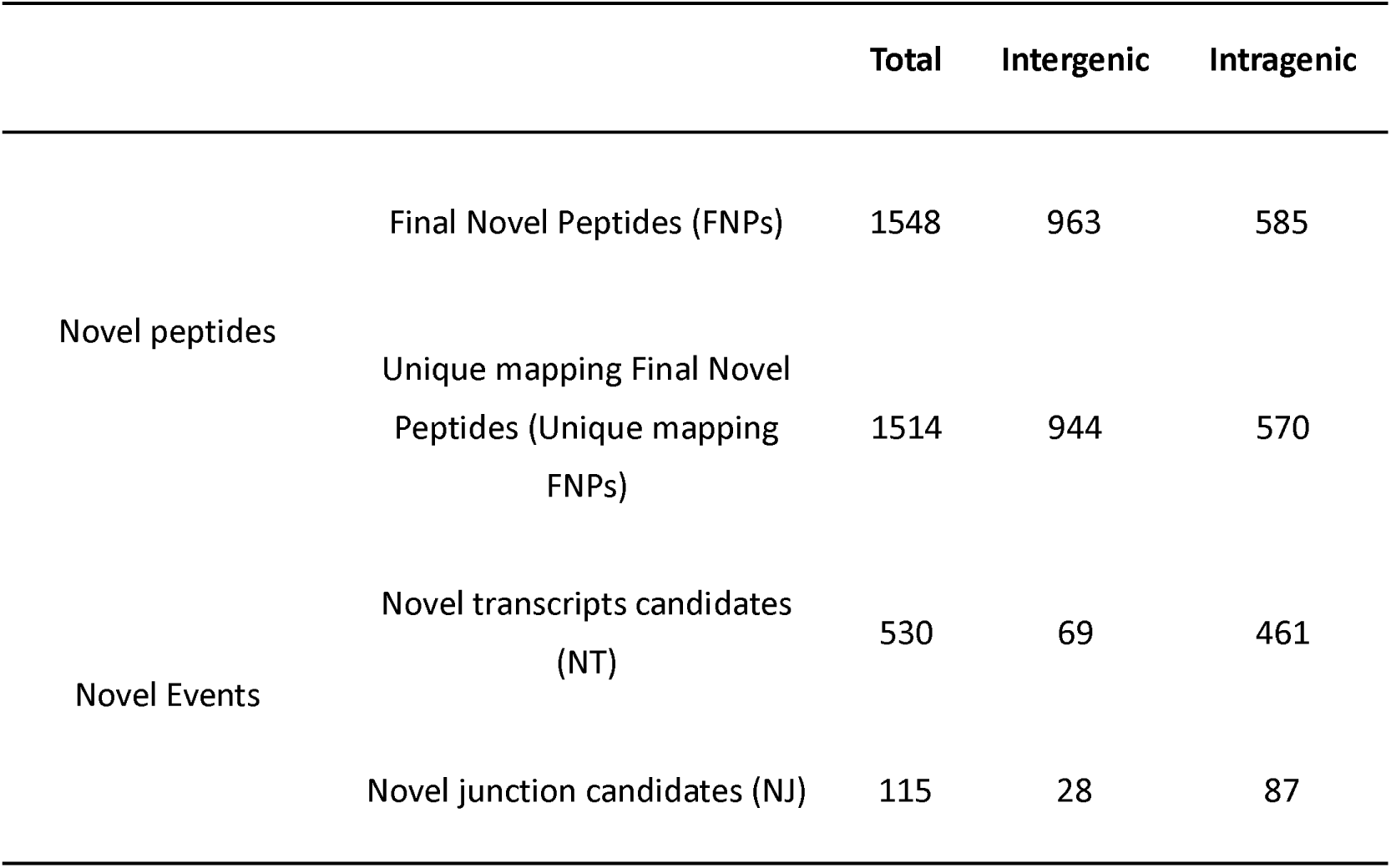
Classification of the novel peptides and novel events based on their type and localization.

|  |  | Total | Intergenic | Intragenic |
| --- | --- | --- | --- | --- |
| Novel peptides | Final Novel Peptides (FNPs) | 1548 | 963 | 585 |
|  | Unique mapping Final Novel Peptides (Unique mapping FNPs) | 1514 | 944 | 570 |
| Novel Events | Novel transcripts candidates (NT) | 530 | 69 | 461 |
|  | Novel junction candidates (NJ) | 115 | 28 | 87 |

For the sake of tracing all the FNPs back to rice genome, we first clustered the NEs based on their genomic loci, because the NJs and NTs generated from different algorithms might have some overlapping sites or regions on the genome. We then localised all the FNPs to these correspondingly clustered NE regions. A total 686 clusters are found on the genome, of which 645 contain at least one unique mapping FNP. The 645 clusters are further divided to two groups according to whether a cluster contains at least one novel transcript or not, the former termed as novel transcript clusters (NT clusters) totalling 530 clusters, and the others denoted as novel junction clusters (NJ clusters) totalling 115 clusters. In the NT clusters, the novel transcripts suggest the putative gene models of the region, and the novel junction are considered as a complementary part of the gene model. If an NT cluster is mapped to an intergenic region, it would be considered as a novel gene candidate, while if it is overlapped with an intragenic region, it would be denoted as a complementary to a known gene. Once the 530 NT clusters were mapped onto IRGSP-1.0.30, 69 NT clusters were mapped in the intergenic regions, while the other 461 are matched in the intragenic region. The NJ clusters (i.e. with only novel junctions) suggest incomplete gene model, indicated as putative splicing sites of an underlying model. Similarly, they were mapped onto IRGSP-1.0.30, as well, resulting in 28 and 87 NJ clusters localized in the intergenic and intragenic region, respectively. The details of the NT clusters and NJ clusters mapped to rice genome are summarized in Table 3, and the two typical examples of NT and NJ cluster are depicted in Figure 4A and B.

**Figure 4.**
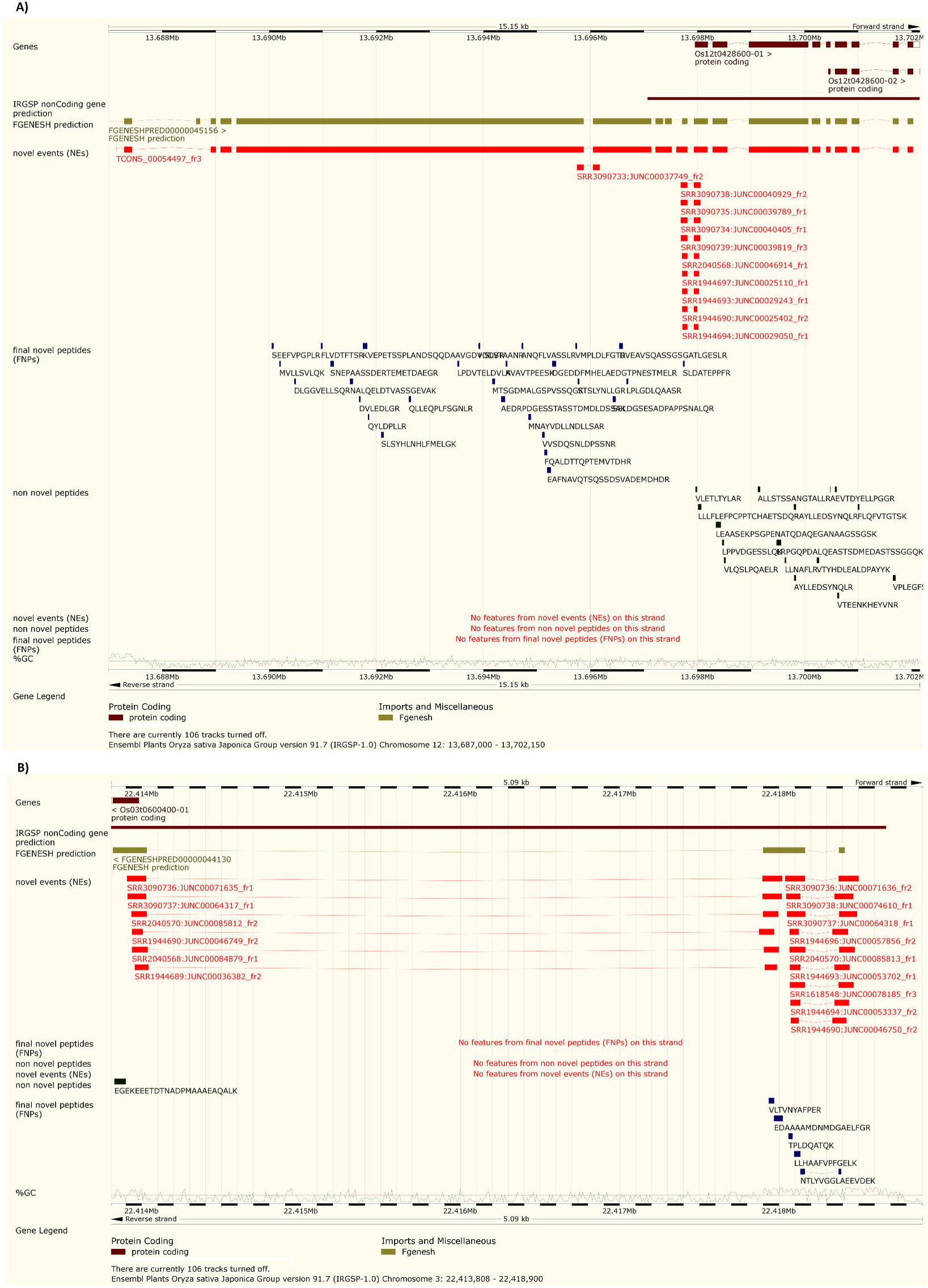
Genomic landscapes of the two typical NEs found in this study. **A)** Both evidence types, transcriptomics and proteomics, support the encoding regions located upstream of the of the 0sl2t0428600 gene. A screen shot is obtained from Gramene, showing the genomic information of position 13687144-13702026 on rice chromosome 12. The track names on the illustrate evidence from different sources, including predicted genes, NEs and novel peptides. On the right, the genomic landscape for the gene 0sl2t0428600 from a theoretical prediction and official annotation, and the location of the transcripts and peptides identified on the same gene based on the official annotation and the NEs. **B)** A screen shot is obtained from Gramene, showing the genomics information of 22413908-22418499 on rice chromosome 3. The three novel peptides and especially the junction peptide (NTLYVGGLAEEVDEK) suggest a novel encoding region and confirmed its junction site.

### Evaluation towards the NEs with the evidence of FNPs

After the FNPs and their corresponding NEs had been clustered, we compared the NEs supported by FNPs with coding and noncoding genes from IRGSP-1.0.30. First, the length statistics of the NT clusters was estimated, i.e. selecting the longest transcript from all the same clustered transcripts. As shown in a violin-plot of transcript length in Figure 5A, the length distribution of the coding and noncoding transcripts in IRGSP-1.0.30 is substantially different from each other. The distribution of NEs is also similar to that of coding transcripts but very distinct from noncoding transcripts. Second, the junction sites (nucleotide pairs on either site of the site junction) mapped from FNPs were compared with known junction sites from IRGSP-1.0.30 for coding and noncoding genes (Supplementary Figure 6). The distributions of junction site distributions for NJs was near identical to junctions in the official annotation, and substantially different to the distribution of splice junction sites for noncoding junctions.

**Figure 5.**
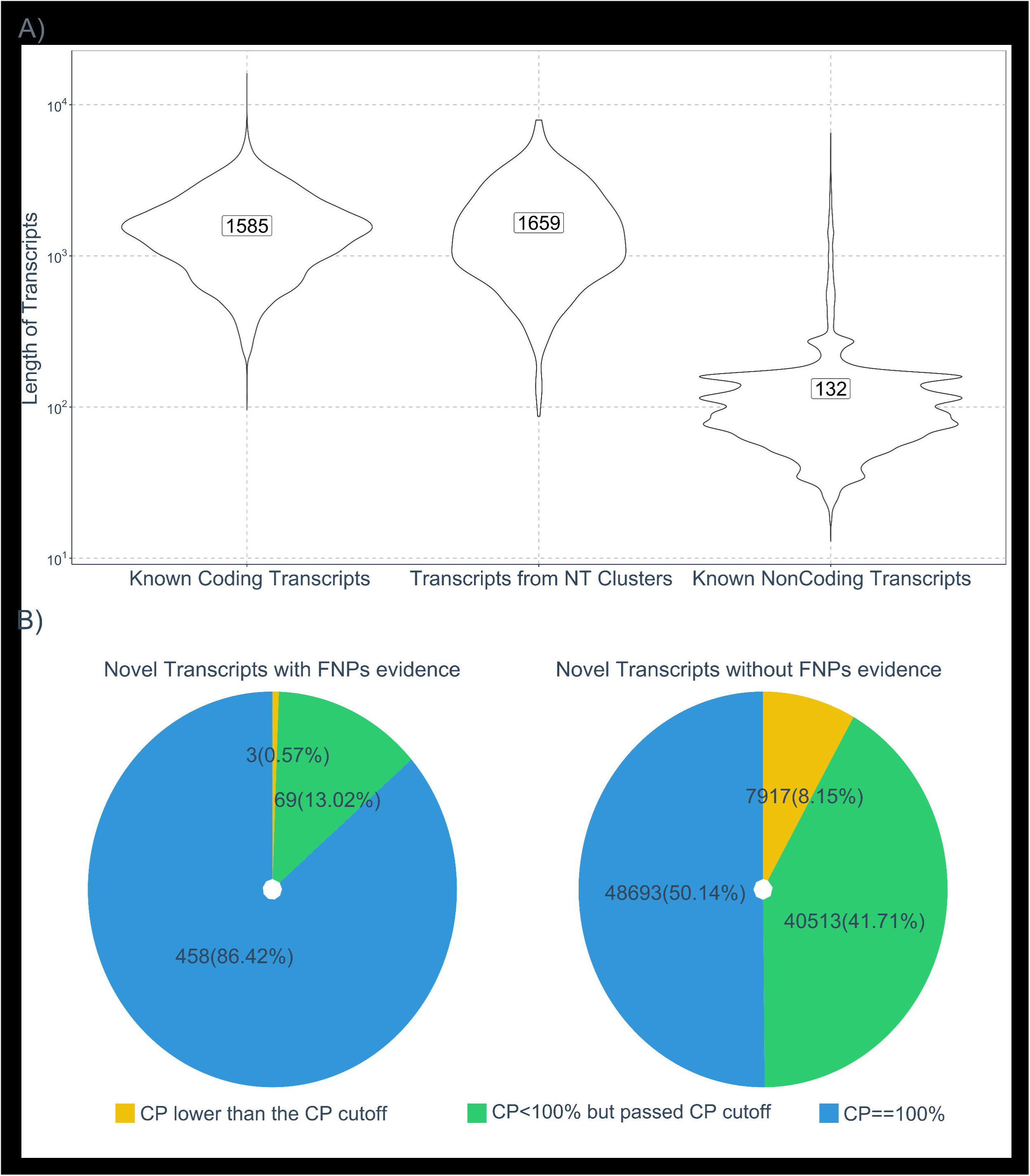
Evaluation to the NEs with the evidence of FNPs. **A)** The length distribution of novel transcripts from NEs with evidence of FNPs (left) compared with coding (middle) and noncoding transcripts (right) from IRGSP-1.0.30. **B)** The coding potential (CP) of novel transcripts from NEs with evidence of FNPs and without evidence of FNPs.

The CPAT pipeline is a well-accepted method for estimating the coding potential of transcripts. We used CPAT to evaluate all the NTs for their coding potential. As shown in Figure 5B, the evaluation suggests that 86.42% of the NTs supported by FNPs (530) have fully confident CPs (scored as 100% by CP), whereas less than 1% of that NTs have CPs below the CP coding threshold (see supplementary Figures 4 and 5 for training data). The analysis of NTs without evidence from FNPs (97123), shows that only 50% of transcripts are scored as coding with full confidence (CP=100%). We thus demonstrate a strong enrichment for transcripts supported by FNPs to be coding.

### Support for novel peptides in other annotations

We performed a BLASTp (“BLASTp short”) search of novel peptides against proteomes from Ensembl plants to detect if these regions had been predicted as genes in other species or varieties of rice (Supplementary File 1, tab “BLASTp final novel peptides”). Out of 1,584 FNPs, only 321 are not matched to other species following stringent thresholding, with 1,263 having at least one hit to another species, indicating that the majority of FNPs are annotated at least once in another species. A heat map and distribution of hits to the species is displayed in Figure 6. The novel peptides could mainly be found in *poaceae* plant (grasses), especially varieties of rice *oryza* and related species. As the peptides’ length increase, the matches to distant species are seen less often, presumably due to the stringent nature of the thresholding we are applying (only one mismatch or deletion allowed).

**Figure 6.**
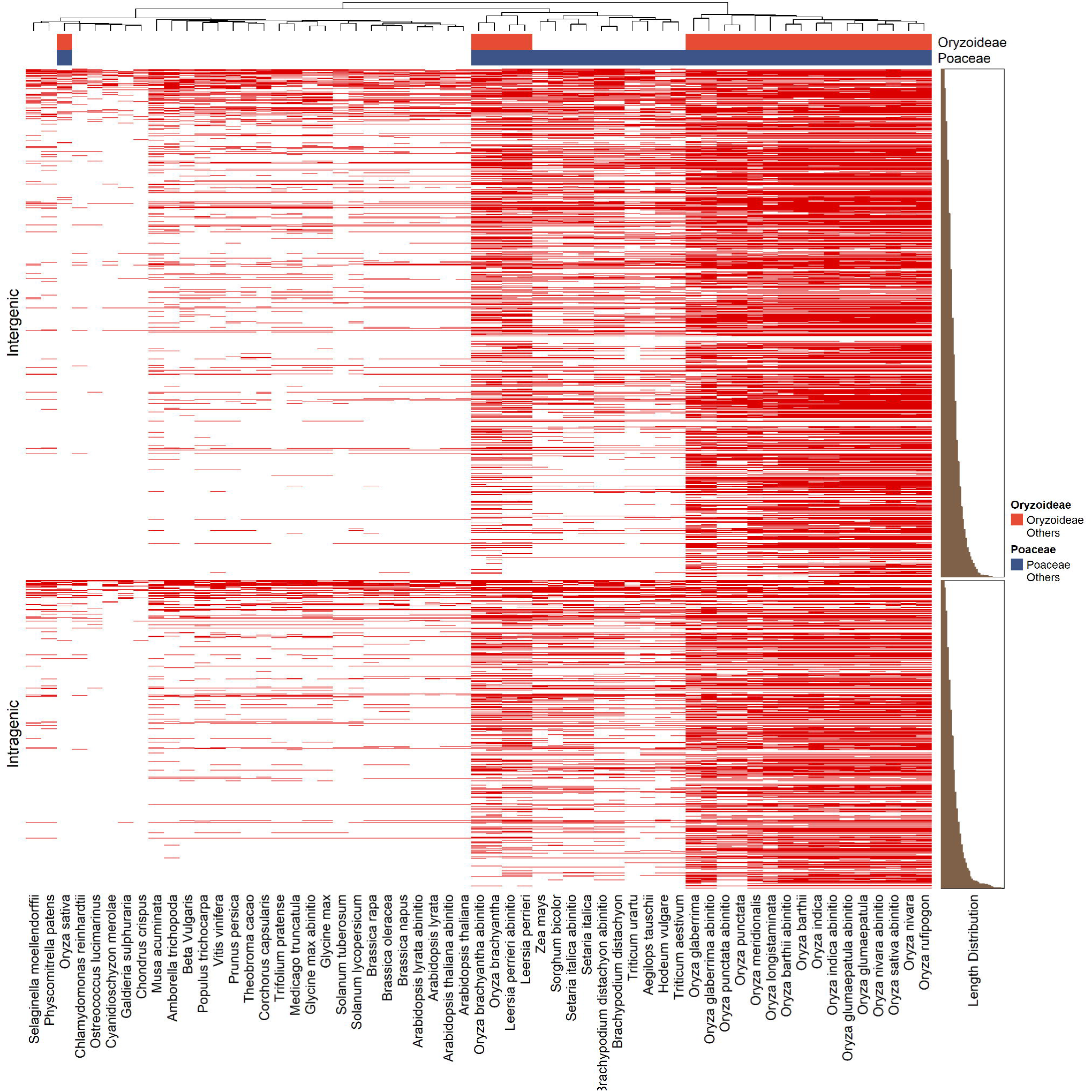
Identification of novel peptides in annotations from other plants. The heatmap represents the hierarchical analysis with the novel peptides against the proteins encoded by the 44 plant genomes from Ensembl (red = positive match, white = no match from BLASTp). The novel peptides are divided to two groups, intergenic (upper panel) and intragenic (lower panel), and are ranked by the peptide lengths for hierarchical analysis.

To further support that NEs supported by FNPs are protein-coding, we performed InterProScan analysis on the translation sequences of 101 intergenic transcripts from 69 NT clusters (full details in Supplementary File 1, tab “InterProScan summary”. Out of 101 sequences, 43 sequences had a significant (e-value < E-05) match to an existing protein domain or family signature, indicating strong evidence that these are indeed missed genes in the annotation. We found evidence for new genes with functions including DNA repair, hydrolase, urease, thiolase-like, ATP synthase and proteinase inhibitor.

### Visualization of results and public availability

To support on-going annotation efforts, we have made results available as permanent Track Hubs. In Supplementary File 1 (tab “Final Novel Peptides”), there is a link from each cluster supported by one or more novel peptides to a visualization of the corresponding region viewed on the Ensembl genome. There are instructions in the Supplementary material (Supplementary Figure 3) for how to configure the tracks to make them visible, which must be followed first.

### Conclusions

We have performed a rigorous proteogenomics analysis on rice, to support future efforts to improve the annotation of the genome. Our results have been openly released in standard formats, which can be easily viewed directly as tracks on the genome browsers. As well as providing confirmatory evidence for over 8000 genes as being protein coding, we have strong evidence that the annotations can be improved for >600 loci. A total of 101 loci were identified in intergenic regions with strong peptide support, of which 43 had a positive prediction of a functional domain, indicating new genes that can be added to the rice genome. Since the results are made persistently available as genomic tracks, we anticipate that future curation efforts will use these datasets for determining the protein-coding gene set of rice.

